# Macrovascular blood flow and microvascular cerebrovascular reactivity are regionally coupled in adolescence

**DOI:** 10.1101/2024.04.26.590312

**Authors:** Kristina M. Zvolanek, Jackson E. Moore, Kelly Jarvis, Sarah J. Moum, Molly G. Bright

## Abstract

Cerebrovascular imaging assessments are particularly challenging in adolescent cohorts, where not all modalities are appropriate, and rapid brain maturation alters hemodynamics at both macro- and microvascular scales. In a preliminary sample of healthy adolescents (n=12, 8-25 years), we investigated relationships between 4D flow MRI-derived blood velocity and blood flow in bilateral anterior, middle, and posterior cerebral arteries and BOLD cerebrovascular reactivity in associated vascular territories. As hypothesized, higher velocities in large arteries are associated with an earlier response to a vasodilatory stimulus (cerebrovascular reactivity delay) in the downstream territory. Higher blood flow through these arteries is associated with a larger BOLD response to a vasodilatory stimulus (cerebrovascular reactivity amplitude) in the associated territory. These trends are consistent in a case study of adult moyamoya disease. In our small adolescent cohort, macrovascular-microvascular relationships for velocity/delay and flow/CVR change with age, though underlying mechanisms are unclear. Our work emphasizes the need to better characterize this key stage of human brain development, when cerebrovascular hemodynamics are changing, and standard imaging methods offer limited insight into these processes. We provide important normative data for future comparisons in pathology, where combining macro- and microvascular assessments may better help us prevent, stratify, and treat cerebrovascular disease.

## Introduction

Cerebrovascular disease is a common risk factor for stroke, a leading cause of disability and mortality across the lifespan.^1–3^ The severity of cerebrovascular disease and corresponding stroke risk are commonly evaluated using diagnostic imaging techniques to visualize cerebral blood vessels, including ultrasound, digital subtraction angiography (DSA), CT angiography (CTA), and magnetic resonance angiography (MRA).^4–6^ These assessments focus on *macrovascular* health, such as the presence of stenosis in large cerebral arteries stemming from the Circle of Willis. However, these imaging assessments are limited because they do not provide information on how macrovascular pathology translates to *microvascular* impairment (small blood vessels at the tissue-level).^4^ For example, occlusion of a large vessel may increase vascular resistance to flow, but downstream vessels (e.g., pial arterioles) can compensate by dilating to reduce overall resistance, resulting in minimal risk to actual tissue.^7,8^ In other cases, microvascular compensatory strategies may be insufficient to maintain cerebral blood flow, increasing the risk for ischemia and necessitating more aggressive interventions.^9^ Understanding the interplay of macrovascular and microvascular hemodynamics would refine our understanding of many cerebrovascular diseases (e.g., stroke, moyamoya, intracranial atherosclerosis) and help guide intervention strategies.

Developing novel cerebrovascular imaging assessments is especially important for adolescents; 15% of all ischemic strokes occur in adolescents and young adults,^10^ but pediatric-specific safety and quality considerations (e.g., motion, radiation exposure, contrast administration, dental hardware) may limit the utility of certain imaging modalities in this population. For example, head motion is higher in children^11,12^, which generally reduces image quality. While CT and PET are frequently used in adults, there is increased concern for exposing younger patients to ionizing radiation and injectable contrast agents.^13–15^ Angiography approaches such as CTA and DSA require catheter insertion via carotid or femoral arteries, an invasive procedure that requires sedation and is further complicated by the smaller caliber of pediatric vessels. Additionally, contrast-based methods cannot be readily repeated within a scan session to obtain multiple hemodynamic measurements. Ultrasound or MRI are typically preferred in pediatric cases, but they also have drawbacks. For example, transcranial Doppler ultrasound (TCD) is commonly used to evaluate stroke risk in sickle cell disease,^16^ but regular TCD examinations often stop at age 16, based on recommendations from the original STOP trial^17^ and poor insonation windows as the skull thickens with age. Thus, many adolescents with sickle cell disease do not receive screening beyond age 16, despite their stroke risk continuing into adulthood. Even when they do, TCD does not provide direct information about microvascular health or tissue-level infarcts^18–20^ which are a prevalent problem in sickle cell disease^21^ and linked to significantly increased stroke risk.^22^

Although MRI offers a non-invasive, versatile alternative to evaluate cerebrovascular health, it is less frequently used in pediatric settings due to longer exam times, which can lead to an increased likelihood of motion artifacts and sometimes necessitate sedation. Nevertheless, standard 2D phase contrast MRI can provide helpful information by directly measuring blood flow velocity but has limitations such as individual placement of imaging planes and single-direction velocity encoding.^23,24^ Methods such as Noninvasive Optimal Vessel Analysis (NOVA)^25,26^ may help to automate and speed up the positioning and acquisition of these 2D phase contrast scans. 4D flow MRI (time-resolved, 3-directional velocity encoding) is another extension of standard 2D phase contrast, allowing full coverage of all major cerebral arteries within one acquisition and retrospective velocity and flow quantification at any position within the imaging volume.^27–29^ Furthermore, 4D flow MRI is gaining clinical application and does not rely on an accessible insonation window like TCD, making it suitable for the complete adolescent age range (approximately 10 to 24 years old).^30^

Additionally, the developing brain undergoes significant neural and vascular remodeling, which increases hemodynamic variability and may alter the relationship between macrovascular and microvascular hemodynamics throughout adolescence.^31–34^ Many hemodynamic parameters, including cerebral blood flow,^35–37^ vascular reactivity,^38^ and flow velocity,^27,39^ increase throughout childhood before declining to adult values in adolescence. Macrovascular arterial pathology is a poor indicator of stroke outcomes in pediatrics,^15^ potentially due to denser pial collateral networks in younger individuals, which may maintain perfusion despite occlusion of a large vessel.^5,14,31^ However, cerebrovascular maturation remains poorly understood^40^ and many adult imaging sequences do not consider the changing composition of the developing brain^41^. In this age range demonstrating rapid developmental changes, it is unclear whether macrovascular or microvascular pathology (or both) are more important for evaluating stroke risk. Furthermore, the expected relationship between macro- and micro-vascular hemodynamics in typical development is unclear.^31,40^ More sophisticated imaging methods are necessary to understand the relationship between macrovascular and microvascular hemodynamics in adolescents and to identify key indicators of cerebrovascular impairment.^4^

Emerging MRI techniques show promise for addressing this need by providing multi-parametric, quantitative evaluation of hemodynamics at different spatial scales without the need for ionizing radiation or contrast. In this study, we examined a novel combination of advanced cerebrovascular MRI techniques to assess the normative relationship between macrovascular and microvascular hemodynamics in healthy adolescents (Figure 1). We acquired 4D flow MRI to obtain blood velocity and blood flow within large arteries originating in the Circle of Willis.^42^ To assess microvascular function, we used cerebrovascular reactivity (CVR), which reflects the health of small vessels regulating blood supply to tissue. CVR is defined as the local blood flow response to a vasoactive stimulus and represents the ability of cerebral arterioles to dilate or constrict.^43–45^ Two aspects of the CVR response are important indicators of vascular health: 1) CVR amplitude, the change in local blood flow per unit stimulus, and 2) CVR delay, when the local blood flow response happens relative to the stimulus timing. CVR has been successfully measured in younger populations using blood-oxygenation-level-dependent (BOLD) functional MRI to measure the blood flow response to increased arterial carbon dioxide during a breath-hold task.^46–49^ Here, CVR amplitude is reported in standard units of %BOLD signal change per mmHg end-tidal CO_2_ (end-tidal gas measurements are a common non-invasive surrogate for arterial CO_2_ measurements). CVR delay, in units of seconds, can be reported in either absolute or relative terms. Absolute CVR delay represents the transit time for hypercapnic blood to travel from the lungs to the heart to the brain, additional transit to a specific brain region, local hemodynamic response dynamics, and their subsequent influence on the venous-weighted BOLD signal.^50–56^ We report CVR delay relative to the gray matter median delay, which removes the effect of transit time to the brain, emphasizing local transit and vasodilatory effects. We report CVR delay relative to the gray matter median delay, which removes the effect of the stimulus duration and transit time to the brain, emphasizing local transit vasodilatory effects. Both CVR amplitude and delay are sensitive to various pathologies, including stroke,^57^ intracranial stenosis,^56,58^ multiple sclerosis,^59^ moyamoya disease,^54,60^ and sickle cell disease.^61,62^

**Figure 1.**
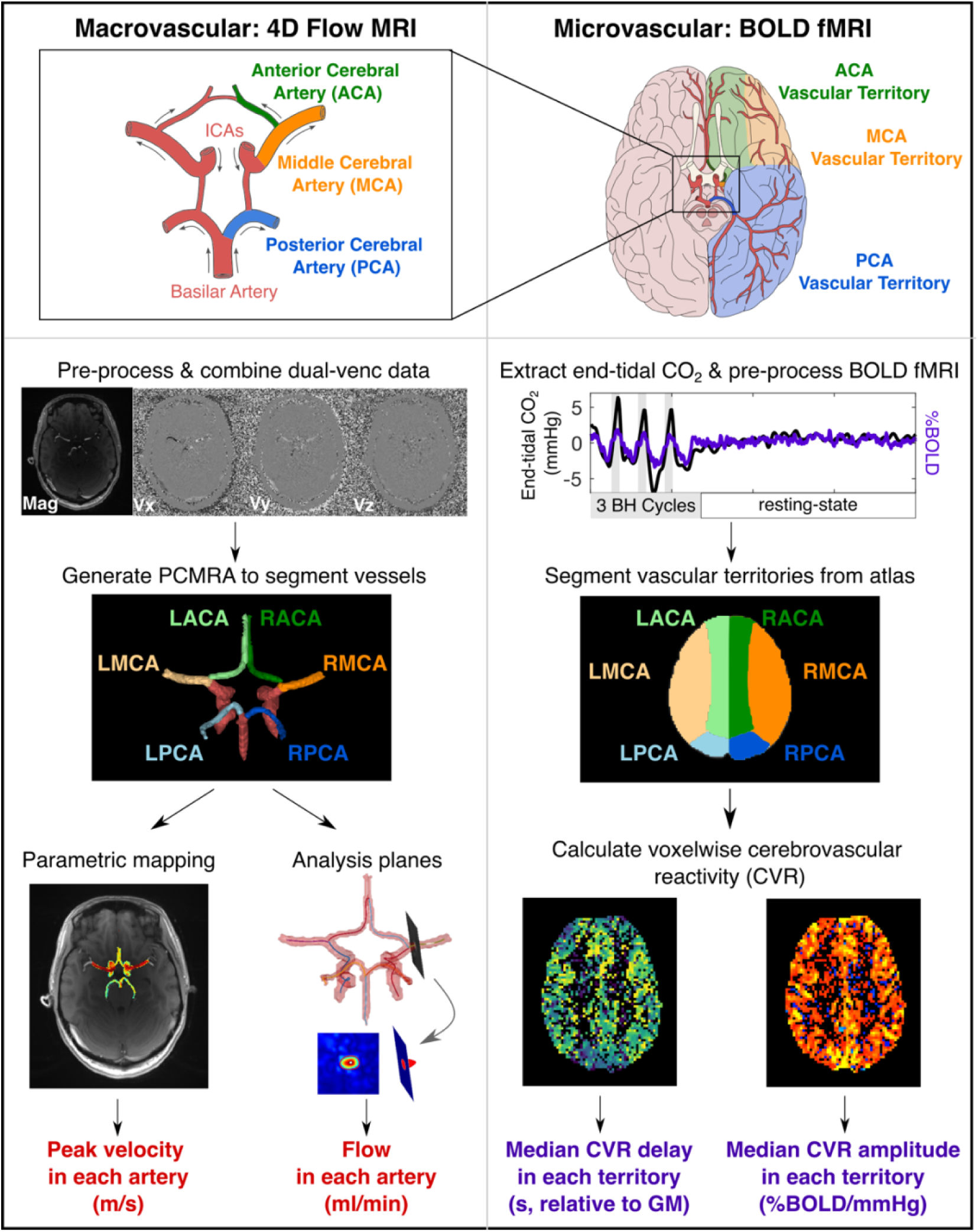
Illustration of the macrovascular and microvascular hemodynamic metrics assessed using MRI in this study. Macrovascular flow in left (L) and right (R) anterior, middle, and posterior cerebral arteries (ACA, MCA, PCA, respectively) was quantified with a single 4D flow MRI acquisition. 4D flow magnitude data (mag) and velocity in 3-directions (Vx, Vy, Vz) were pre-processed to generate a phase-contrast magnetic resonance angiogram (PCMRA). Peak velocity and flow were then extracted from each artery. Cerebrovascular reactivity (CVR) in the associated vascular territories was measured with whole-brain blood-oxygenation-level-dependent functional MRI (BOLD fMRI). During this scan, participants performed a hypercapnic breathing challenge and end-tidal CO_2_ was recorded. Median CVR amplitude and delay were computed within the vascular territories of these arteries.

In a healthy cohort spanning the adolescent age range, we investigated two regional relationships: 1) macrovascular blood velocity and downstream CVR delay, 2) macrovascular blood flow and downstream CVR amplitude. To summarize regional CVR responses, we averaged voxelwise values across a vascular territory, the brain region supplied by each intracranial artery exiting the Circle of Willis. We hypothesized that CVR delay is negatively correlated with baseline blood velocity; in other words, vascular territories supplied by arteries with higher velocity respond earlier to a hypercapnic stimulus due to faster arrival of hypercapnic arterial blood.^58,65–67^ In contrast, we hypothesized that CVR amplitude is positively correlated with macrovascular blood flow, because resistance of large arteries influences downstream microvascular pressure and perfusion,^63,64^ which in turn influence arteriolar dilation capacity.^65–67^ We aimed to pilot these advanced techniques in adolescents and establish a baseline for future comparison in cerebrovascular pathologies where macro- and micro-vascular relationships may be an important indicator of stroke risk. Understanding these relationships in the transition from childhood to adulthood is critical to identify normal developmental variability and to address challenges of current cerebrovascular imaging assessments in this age range.

## Material and Methods

### Participants

This study was approved by the Northwestern University Institutional Review Board (STU00210491). Informed written consent (parental permission and child assent for individuals under 18) was obtained before the MRI session. Participants consisted of 12 healthy adolescents between ages 8 and 25 (7F, 16±6 yrs) with no history of psychiatric or neurological disorders. Individuals younger than 8 were excluded based on prior experience with poor breath-hold performance in this age group.^68^ We extended the age range to 25 to capture the transition from adolescence to adulthood before age-related cerebrovascular decline begins.^69^

To probe how the macrovascular and microvascular relationships might differ in established macrovascular pathology, we included a case study in an adult (M, 33 yrs) with moyamoya arteriopathy of the right MCA. This participant underwent the same MRI acquisitions and analysis pipelines as controls, explained in the following sections.

### Magnetic Resonance Imaging

Participants underwent MRI scanning on a 3T Siemens PrismaFit with a 64-channel head coil. Medical tape was placed across the forehead to minimize head motion.^70^ A 3D axial time-of-flight magnetic resonance angiography was performed to position the 4D flow imaging volume, which was centered on the MCAs (TR = 21.0 ms, TE = 3.42 ms, FA = 18°, resolution = 0.3 × 0.3 × 0.5 mm^3^, slices per slab = 40, slabs = 4). Dual velocity-encoding (*venc*) 4D flow MRI^29^ was collected with TR = 5.9 ms, TE = 3.3 ms, FA = 15°, low-*venc* = 0.8 m/s, high-*venc* = 1.6 m/s, 1 mm isotropic resolution, temporal resolution = 82.6 ms, k-t GRAPPA R = 5, scan time = 10-12 min). A functional MRI scan was collected using a multi-band multi-echo gradient-echo EPI sequence provided by the Center for Magnetic Resonance Research^71,72^ (TR = 1500 ms, TEs = 10.8/28.03/45.26/62.49/79.72 ms, FA = 70°, resolution = 2.5 × 2.5 × 2.0 mm^3^, MB factor = 4, GRAPPA = 2). The number of functional volumes varied based on the breath-hold task. Pairs of single-band reference (SBRef) images with opposite phase-encoding (AP or PA) were collected for each echo time to facilitate functional realignment, masking, and distortion correction. A whole-brain T1-weighted EPI-navigated multi-echo MPRAGE^73^ was acquired (TR/TE1/TE2/TE3 = 2170/1.69/3.55/5.41 ms, TI = 1160 ms, FA = 7°, 1 mm isotropic resolution, scan time = 5 min 12 s). The three echoes were combined using root-mean-square. See Table S1 for additional scan parameters.

### Breath-hold Task

All participants performed an end-expiration breath-hold task during the fMRI acquisition. Nine participants performed a hybrid protocol consisting of 3 breath-hold trials, followed by 7-minutes of rest.^54^ During the rest period, the abstract “Inscapes” video was played to reduce motion.^74^ The other three participants (22-yoM, 24-yoM, 25-yoM) performed 5 breath-hold trials without resting-state. Each breath-hold trial consisted of four paced breathing cycles (6s period), 15s breath-hold, 2s exhalation, and 8s free recovery breathing. Participants were cued with visual, graphical instructions projected through a mirror on the head coil.^68^ Expired CO_2_ concentrations were sampled continuously (1000Hz) via nasal cannula and gas analyzer (ADInstruments). Prior to the scan, subjects practiced the task and were instructed about the importance of exhaling through their nose both before and after the breath-hold period.^49^

### 4D Flow MRI Pre-processing

Dual-*venc* 4D flow MRI data were processed as described previously.^29,75,76^ Background-phase offsets were corrected using Maxwell terms during image reconstruction.^77^ In-house MATLAB software was used to perform eddy-current correction and noise-masking.^78^ High-*venc* data was used to correct for aliasing in the low-*venc* data, resulting in a single anti-aliased dataset.^29^ A time-averaged 3D phase contrast angiogram (PC-MRA) was constructed to depict vessel anatomy. For each subject, the arteries exiting the Circle of Willis, including bilateral anterior cerebral arteries (ACA), middle cerebral arteries (MCA), and posterior cerebral arteries (PCA), were manually segmented from the PC-MRA (Figure 1) using dedicated software (Mimics Innovation Suite, Materialise, Leuven, Belgium). The 3D vessel segmentations were used to mask the 4D flow data.

### Peak Velocity Calculation: 4D Flow Parametric Maps

Peak velocity within each vessel was calculated using an in-house parametric mapping workflow^79^ adapted for intracranial analysis^a^ (Figure 1). Absolute velocity magnitude was determined for each voxel within a 3D segmentation (e.g., within an individual vessel ROI) at each cardiac timepoint. Then, a waveform of velocity over time was generated for each voxel. Time-to-peak, the time at which maximum velocity occurred over the cardiac cycle, was identified for each voxel and averaged within the vessel ROI. A 3D array of maximum velocities in the ROI was computed by finding the maximum velocity in a systolic window consisting of three cardiac timepoints centered at the average time-to-peak. Peak velocity was then calculated as the 97th percentile velocity magnitude within the ROI. Upper percentile velocity values were removed because they are known to exhibit noise in dual-venc 4D flow data;^29^ the specific threshold was chosen empirically during development.

### Flow Calculation: 4D Flow Plane Analysis

A separate analysis tool was used to calculate total flow within each vessel, as described previously.^75,80,81^ Vessels were identified using a centerline approach, and hemodynamic information was evaluated via analysis planes placed every 1mm along the vessel, perpendicular to the centerline (Figure 1). The vessel lumen boundary was automatically segmented with manual adjustment if needed. At each plane, velocity and net flow were extracted over all cardiac timepoints. Planes with outliers were removed using a median filter (sliding window = 3 timepoints; threshold = 1.5 standard deviations from window median).^76^ Velocity-time and flow-time curves were interpolated to complete the R-R interval and estimate behavior at diastole. For each analysis plane, the interpolated flow curves were integrated to calculate total flow (mL/cardiac cycle). The median flow value over all planes was extracted for each vessel.

### Calculating Average Tissue Perfusion from Flow

Flow values (mL/cardiac cycle) were converted to average tissue perfusion (mL/100g/ min) to account for differences in the heart rate and size of each vascular territory across individuals, and to make direct comparisons with previous literature.^82^ Flow values were multiplied by each subject’s heart rate (cardiac cycles/min) to convert to units of mL/min. A vascular territory atlas^83,84^ was transformed to each subject’s anatomical space (Figure 1). Gray matter and white matter volumes were calculated within each territory based on *fsl_anat* tissue segmentation, then summed to compute total tissue volume (mm^3^). Tissue density of 1.06 g/cm^3^ was assumed to estimate the mass of tissue in each territory.^85^ Flow values were divided by this territory-specific tissue mass to calculate average tissue perfusion.

### Functional MRI Pre-processing

MRI pre-processing was performed with a series of custom scripts combining FSL,^86^ AFNI,^87^ and tedana^88^ commands. This pipeline has been described in detail^89^ and code is publicly available at https://github.com/BrightLab-ANVIL/PreProc_BRAIN. See Supplementary Material for details.

### CVR Amplitude and Delay Estimation

End-tidal CO_2_ (P_ET_CO_2_) peaks were identified with a peak detection algorithm in MATLAB (MathWorks, Natick, MA), manually inspected, linearly interpolated to create P_ET_CO_2_ timeseries, and convolved with the canonical hemodynamic response function to model the breath-hold task (Figure 1). P_ET_CO_2_ was used as the regressor of interest to compute voxelwise estimates of CVR amplitude and delay in a “lagged” general linear model framework with phys2cvr.^53,90^ This method has been described previously.^53,54,91^ Specific analysis choices are detailed in Supplementary Material. Two maps were generated: CVR amplitude (in %BOLD normalized to mmHg P_ET_CO_2_) and CVR delay (in seconds) (Figure 1). Delay maps were centered on the median delay across gray matter voxels. Both CVR amplitude and delay maps were thresholded to remove voxels at or adjacent to boundary conditions.^53^ CVR amplitude and delay maps were normalized to the MNI152 6^th^ generation template (FSL version, 2mm resolution).

### Macro- and Micro-vascular Comparisons

A vascular territory atlas in MNI space^83,84^ was used to define the left and right anterior, middle, and posterior cerebral artery territories (ACA, MCA, PCA, respectively; Figure 1). CVR amplitude and delay maps, t-statistic masks, and gray matter masks were all transformed to MNI space. Median CVR amplitude and delay were calculated in each territory, including only gray matter voxels with positive CVR amplitude and a significant fit (α=0.05) for the end-tidal CO_2_ regressor. White matter voxels were excluded due to lower BOLD SNR in this region.^92^ Two different comparisons were made between macrovascular and microvascular hemodynamics across 6 arteries and their respective territories:

1. Peak velocity & median CVR delay
2. Average tissue perfusion (derived from flow) & median CVR amplitude

For each comparison, a linear mixed effects model was implemented with CVR delay or CVR amplitude as the dependent variable. Peak velocity or average tissue perfusion in the supplying artery were group-mean-centered and included as the fixed effect. Subjects were included as a random effect, allowing for random slopes and intercepts. These estimated parameters (Table 1) were used to create within-subject predictions (Figure 4).

**Table 1.** Linear mixed effects model results for macrovascular and microvascular relationships.

| Dependent Variable: CVR Delay |  |  | Dependent Variable: CVR Amplitude |  |  |
| --- | --- | --- | --- | --- | --- |
| **Fixed Effect:**<br>**Peak Velocity** | Estimate ± SE | t-statistic | **Fixed Effect:**<br>**Avg. Tissue Perfusion** | Estimate ± SE | t-statistic |
| Slope | $-1.055 \pm 0.361$ | -2.925** | Slope | $0.001 \pm 0.0003$ | 3.856*** |
| Intercept | $-0.121 \pm 0.064$ | -1.888 | Intercept | $0.385 \pm 0.026$ | 14.747*** |
| **Random Effect:**<br>**Subject** |  |  | **Random Effect:**<br>**Subject** |  |  |
|  | Variance |  |  | Variance |  |
| Slope | 0.727 |  | Slope | 0.001 |  |
| Intercept | 0.033 |  | Intercept | 0.089 |  |
| Residual | 0.103 |  | Residual | 0.027 |  |

Note: \*p < 0.05, \*\*p < 0.01, \*\*\*p < 0.001

### Left vs. Right Comparisons

In the healthy cohort, we explored whether the macrovascular-microvascular trends were consistent between hemispheres. For each comparison (peak velocity & CVR delay; tissue perfusion & CVR amplitude), we computed two different linear mixed effects models with the framework described above but using data from only left or right hemispheres. These estimated parameters (Tables S2-S3) were used to create within-subject predictions (Figure S1).

### Exploring Trends with Age

This study was not powered to model the effects of age explicitly, but we explored age-related trends in the relationships between macrovascular and microvascular parameters by plotting the predicted slopes for each subject from the linear mixed effects model against age.

## Results

Figure 2 shows macro- and micro-vascular maps and their regional relationships in a representative subject (18-yoF). Peak velocities displayed in the maximum intensity projection map follow expected patterns, with lowest velocities in the PCAs and highest velocities in the MCAs. The CVR delay map shows the timing of the local breath-hold response, relative to the median in gray matter. Within each vascular territory, the distribution of delays is centered near the gray matter median, with later (positive) relative delays in PCA territories and earlier (negative) relative delays in MCA and ACA territories. There is a negative relationship between peak velocity and median CVR delay across the 6 vascular territories.

**Figure 2.**
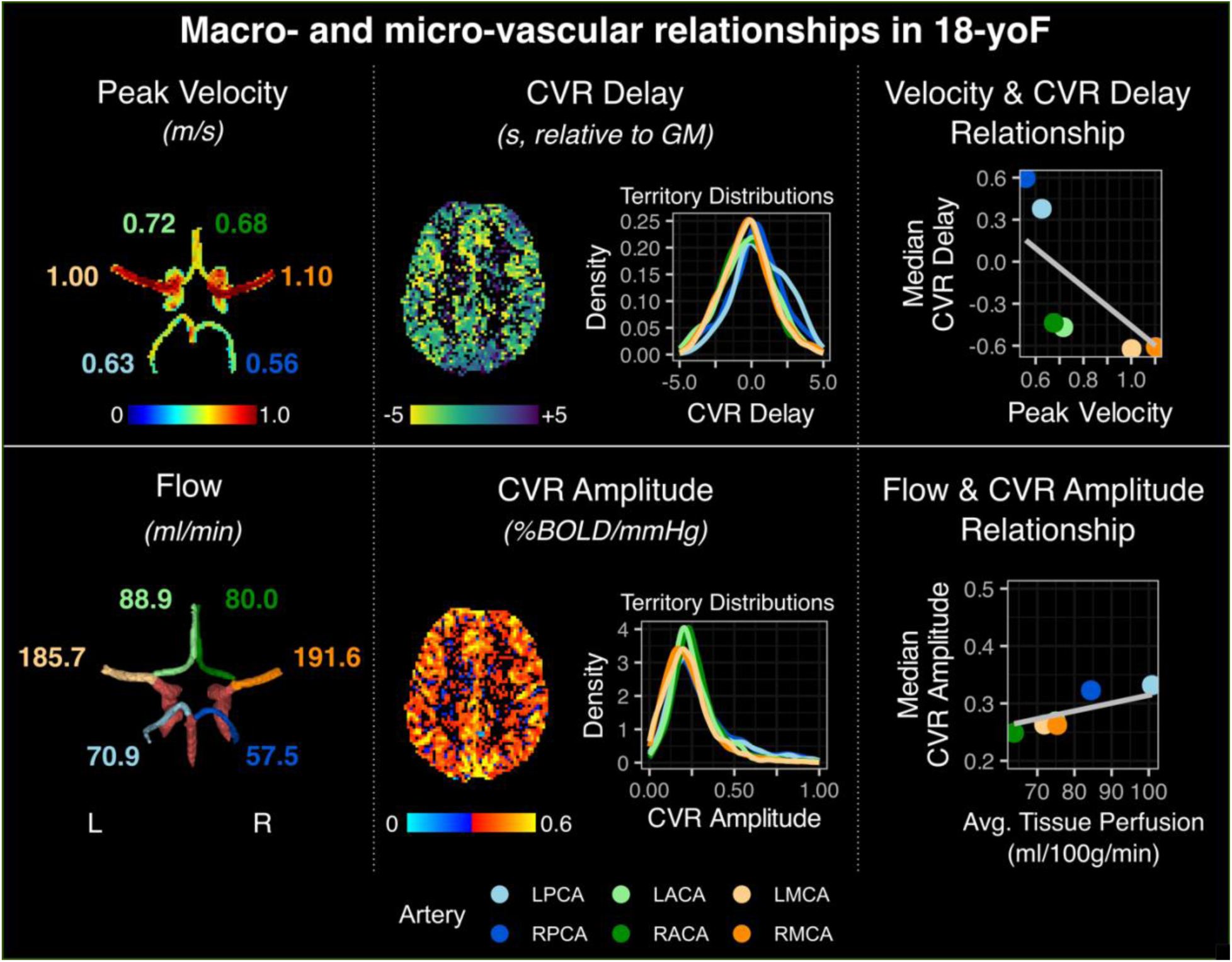
Maps and hemodynamic relationships in a representative adolescent subject. **Top)** A 2D projection of peak velocities throughout the Circle of Willis is displayed, with the peak velocity listed for each artery. A map of cerebrovascular reactivity (CVR) delays (in seconds, relative to the gray matter median) and their distributions within each vascular territory are shown. Relationships between median CVR delay and peak velocity for each vessel are plotted. **Bottom)** Flow values are reported next to each artery. A map of CVR amplitude and the distributions within each vascular territory are shown. Relationships between median CVR amplitude and average tissue perfusion for each vessel are plotted. Note: average tissue perfusion represents flow normalized to territory volume**. Vessel abbreviations:** anterior cerebral artery (ACA), middle cerebral artery (MCA), posterior cerebral artery (PCA).

Flow follows the same trend as peak velocity, with lowest flow in the PCAs and highest flow in the MCAs. However, after accounting for the total volume of tissue in each vascular territory, the trend is reversed—average tissue perfusion is highest in the PCAs. The CVR amplitude map shows the magnitude of the local breath-hold response per unit increase in arterial CO_2_ pressure (in units of millimeters of mercury, mmHg). The median CVR amplitude is highest in the PCA territories and lowest in ACA territories. There is a positive relationship between average tissue perfusion and median CVR amplitude across the 6 vascular territories.

### Peak Velocity & CVR Delay Comparisons

At the group level, the negative trend between peak velocity and CVR delay is consistent; when peak velocity in a supplying artery is higher (as in the MCAs), the downstream vascular territory responds earlier to a breath-hold stimulus. Figure 3 shows the relationship between peak velocity and CVR delay across all subjects and for each subject individually. Linear mixed-effects modeling confirmed this trend (Table 1), with a significant fixed effect of peak velocity on CVR delay. The slope of this relationship is −1.055±0.361, indicating that for each ∼10cm/s increase in peak velocity, CVR delay downstream is ∼0.1s earlier relative to the gray matter median. However, this relationship is variable between subjects, with the subject random effect explaining 88% of residual variance in CVR delay values. The effects of sex and age were not explored in this model due to the small sample size, but trends with age were explored separately.

**Figure 3.**
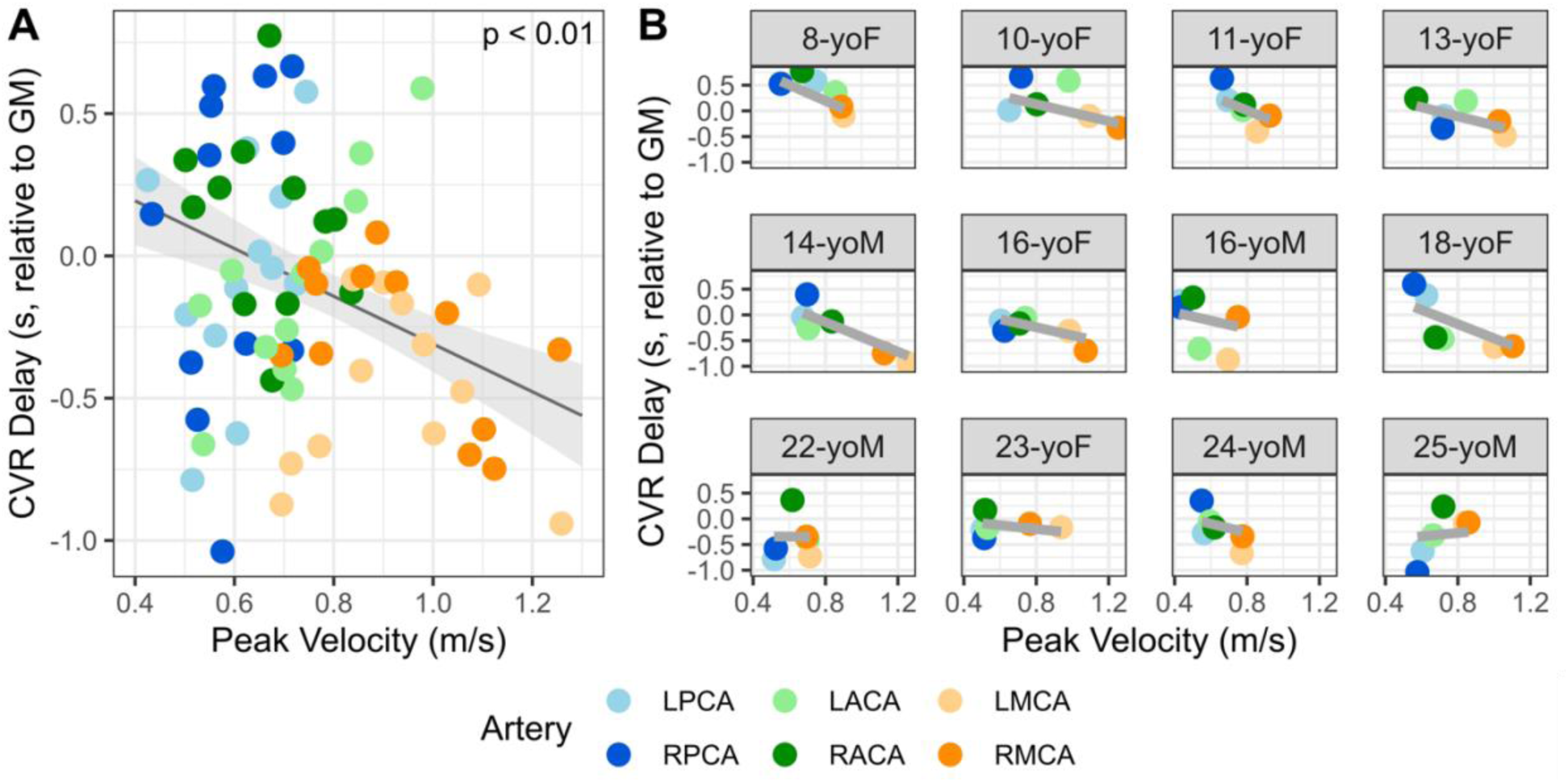
Peak velocity and CVR delay relationships for **A)** the group and **B)** each subject. Each comparison represents peak velocity in an artery and median CVR delay in the associated vascular territory. Linear fits were plotted using estimates from the linear mixed effects model detailed in Table 1. Note that CVR delay is in units of seconds, relative to the median in gray matter for each subject. **Vessel abbreviations:** anterior cerebral artery (ACA), middle cerebral artery (MCA), posterior cerebral artery (PCA).

### Average Tissue Perfusion & CVR Amplitude Comparisons

The positive trend between average tissue perfusion and CVR amplitude is also consistent at the group level; when average tissue perfusion in a supplying artery is higher (as in the PCAs), the downstream territory has a larger BOLD fMRI response to a breath-hold stimulus. Figure 4 shows the relationship between average tissue perfusion and CVR amplitude across all subjects and for each subject individually. Note the importance of considering vascular territory size to this relationship (Figure 4C-F). Without accounting for territory volume, flow (in units of ml/min) is highest through the MCA (Figure 4C). However, the MCAs also have the highest volume of tissue to supply (Figure 4D). After accounting for this fact, the MCAs in fact have the lowest flow per 100g of tissue supplied, while PCAs have the highest (Figure 4E).

**Figure 4.**
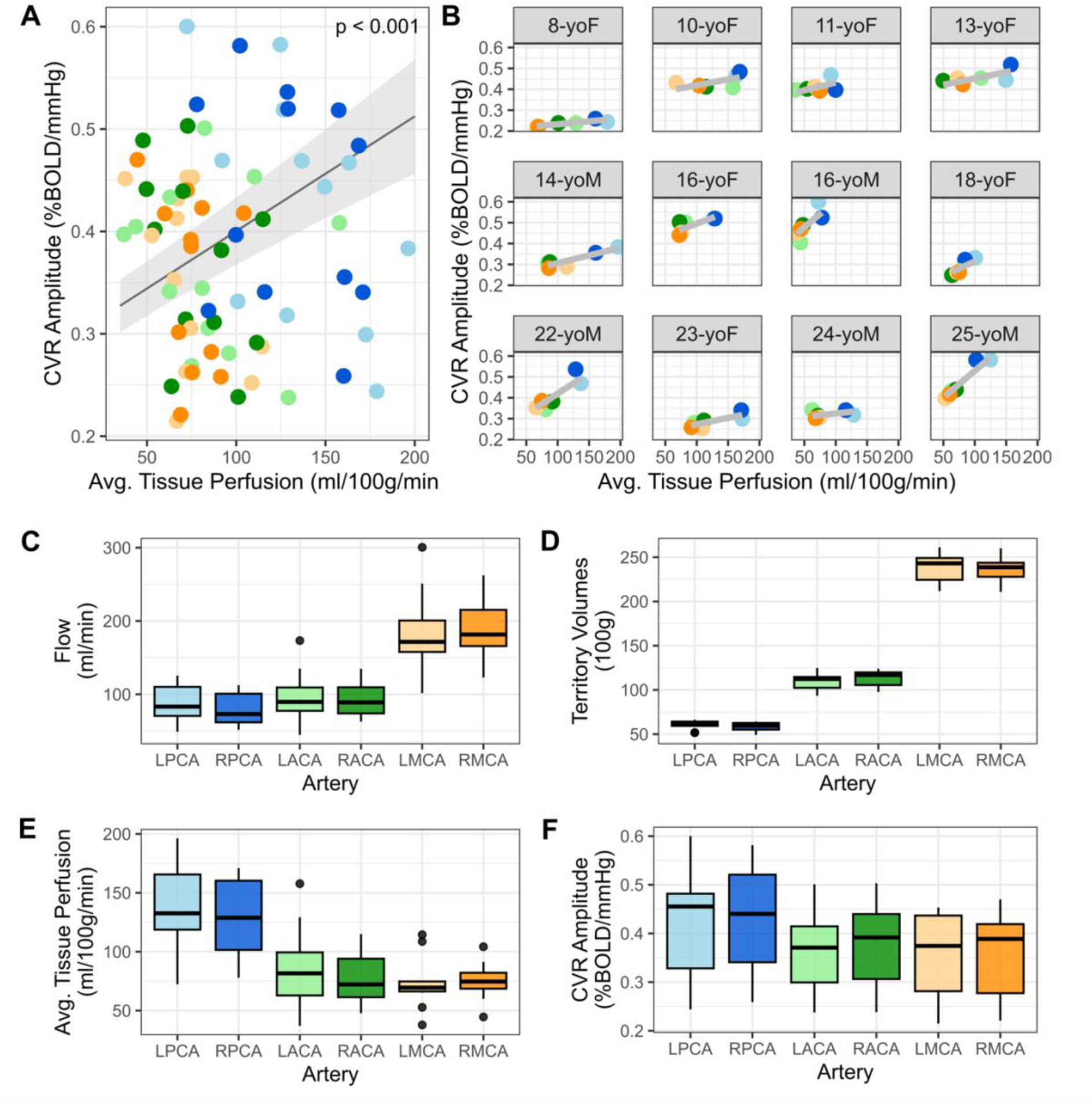
Average tissue perfusion and CVR amplitude relationships for **A)** the group and **B)** each subject. Each comparison represents average tissue perfusion (flow normalized to territory volume) for a given artery and median CVR amplitude in the associated vascular territory. Linear fits were plotted using estimates from the linear mixed effects model detailed in Table 2. The impact of normalizing flow to territory volume is demonstrated by boxplots in panels C-F, which summarize the spread of values across all subjects. **C)** Flow through each artery in standard units of ml/min. **D)** Volume of gray matter and white matter tissue in each territory (in units of 100 grams). **E)** Average tissue perfusion, computed by dividing flow by territory volume. **F)** CVR amplitude in each vascular territory. **Vessel abbreviations:** anterior cerebral artery (ACA), middle cerebral artery (MCA), posterior cerebral artery (PCA).

Linear mixed-effects models confirmed the positive trend (Table 1), with a significant fixed effect of average tissue perfusion on CVR amplitude. Both the slope and intercept of this relationship are significant; the small slope magnitude is due to the difference in magnitude between CVR and perfusion units. However, there is again substantial variability between subjects, with the subject random effect explaining 77% of residual variance in CVR amplitude values. As with CVR delay, trends with age were explored separately from the linear-mixed effects model.

### Left vs. Right Comparisons

The macrovascular-microvascular trends are relatively consistent between left and right hemispheres, with group-level slope estimates similar to those computed when combining hemispheres (Tables S2-S3). This relationship is variable between subjects (Figure S1) but generally follows similar trends.

### Trends with Age

Cerebral hemodynamics are known to change throughout adolescence (ages 10-24),^27,35–39^ so we explored age-related trends in macrovascular-microvascular relationships in our small cohort that spans this developmental timeframe. The small sample size of this initial study precludes explicitly incorporating age in the linear models above, so we instead plotted changes in the linear model slopes with age.

Overall, the relationship between CVR delay and peak velocity is reduced with age (Figure 5A), indicated by the smaller negative slopes in older adolescents. In contrast, the relationship between CVR amplitude and average tissue perfusion strengthens with age (Figure 5B). This trend was not statistically significant (R^2^=0.21, p=0.13) but improved after removing the 16-yo outlier (R^2^=0.32, p=0.07). Thus, age-related changes in macrovascular and microvascular relationships are evident for both timing and flow parameters.

**Figure 5.**
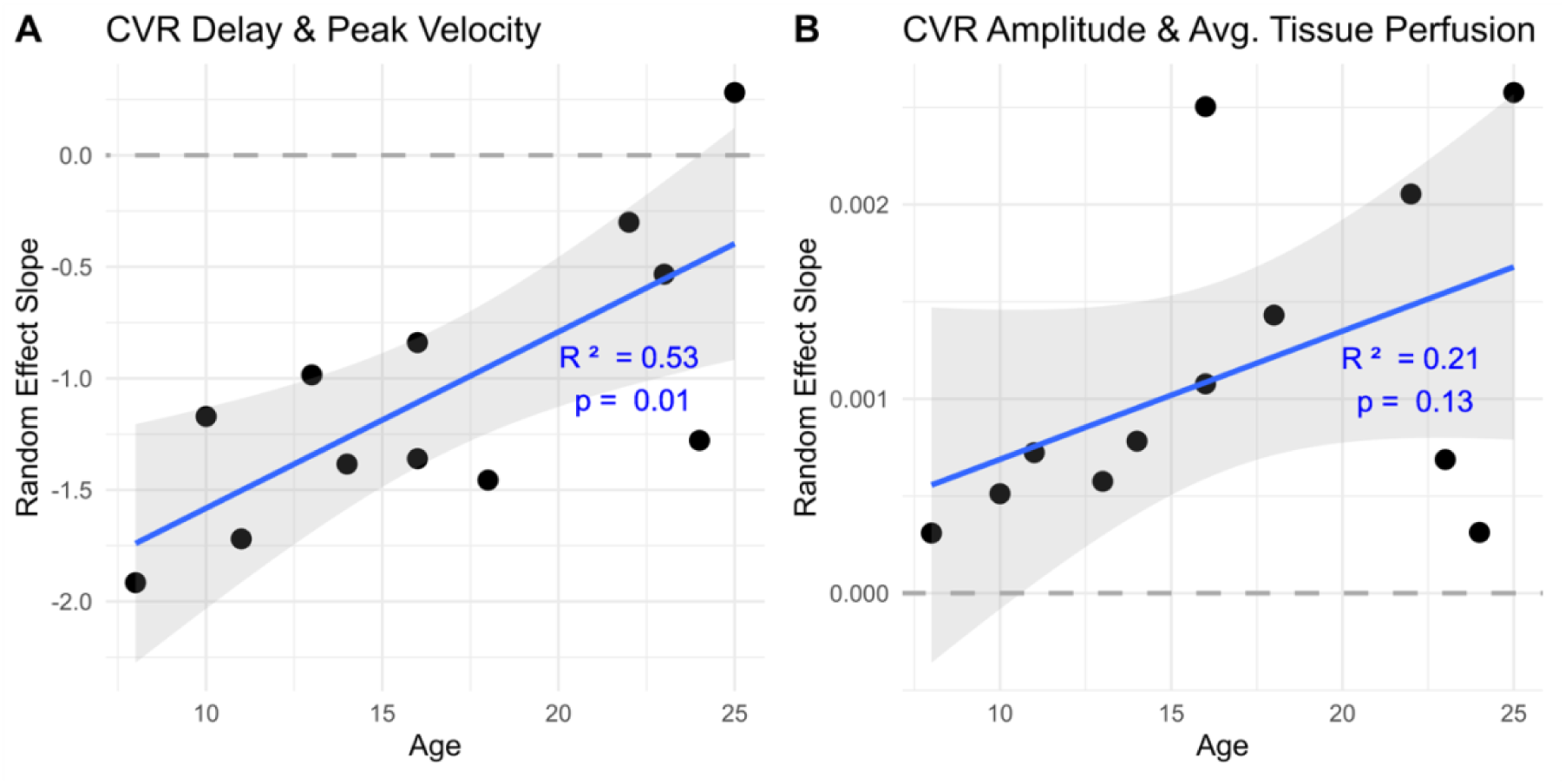
Trends in the slope of macro- and micro-vascular hemodynamic relationships with age. **A)** Slopes between relative cerebrovascular reactivity (CVR) delay and peak velocity decrease with age. **B)** Slopes between CVR amplitude and average tissue perfusion increase with age. A dashed reference line with slope = 0 is shown in each plot.

### Moyamoya Case Study

A board-certified neuroradiologist (SJM) confirmed moyamoya arteriopathy involving the most proximal M1 segment of the right MCA in a 33-year-old male. The time-of-flight image in Figure 6 illustrates how the vessel abruptly ends at this point. Adjacent collaterals, characteristic of moyamoya pathology, are also visible at higher resolution. Unsurprisingly, peak velocity is substantially lower in the RMCA compared to the LMCA. CVR delay in the RMCA territory is also 5 to 10 seconds longer relative to the rest of the brain, consistent with our previous reports in this individual.^54^ However, the CVR delay map suggests only a portion of the RMCA territory is affected by longer CVR delays, likely due to collateral flow. Flow is also substantially lower in the RMCA, and there appears to be compensation through the RACA and RPCA, evidenced by higher flow values in these vessels. CVR amplitude in the RMCA territory is relatively preserved, though the region of negative amplitudes (blue voxels) may be indicative of the vascular steal phenomenon.^93^ Relationships between macro-vascular and micro-vascular hemodynamics follow the same trends as controls (Figures 3-4), with an obvious outlier in the RMCA territory for the velocity and CVR delay relationship.

**Figure 6.**
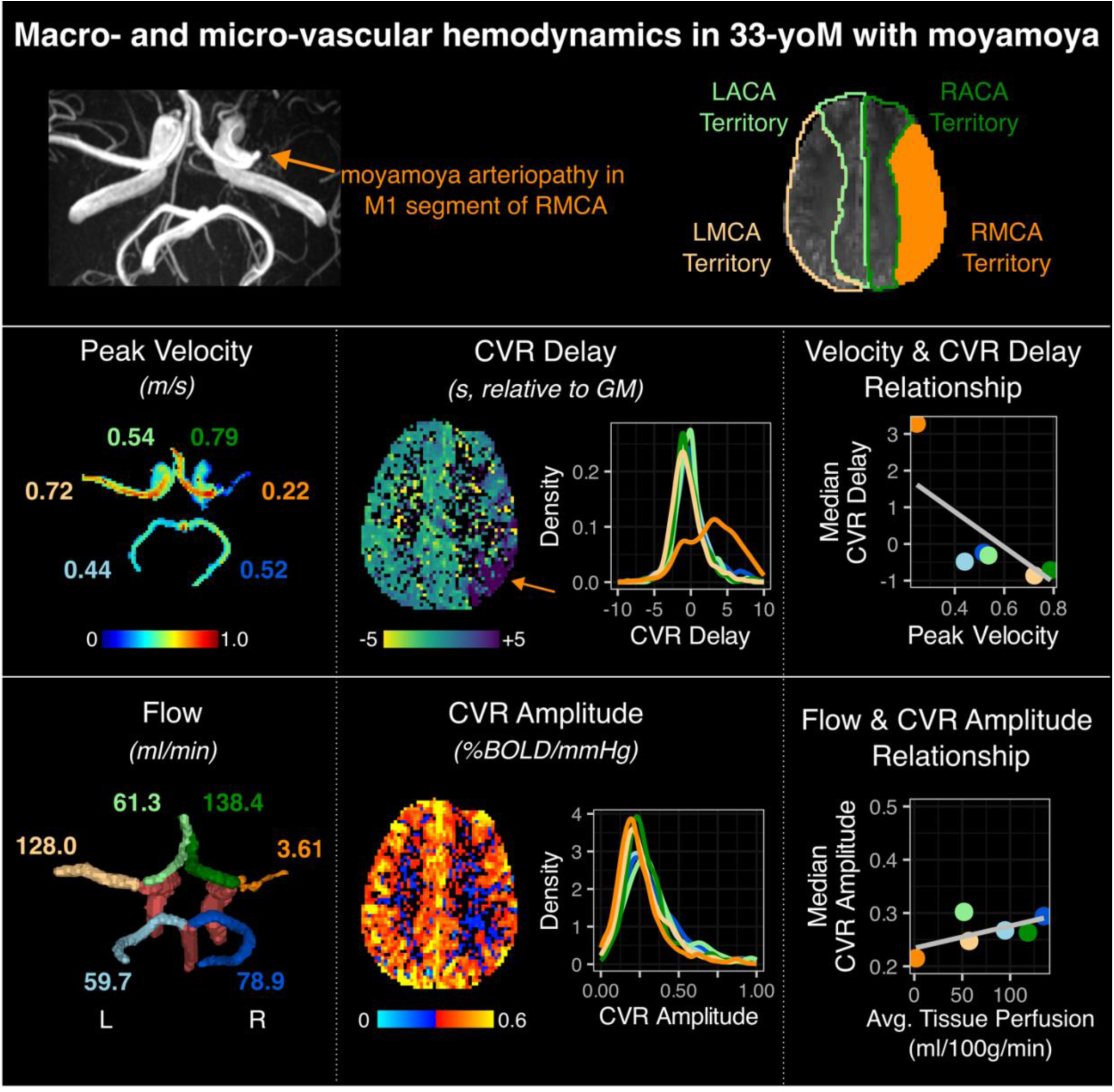
Maps and hemodynamic relationships in an adult male with moyamoya arteriopathy of the right middle cerebral artery (RMCA). Vessel abbreviations and color schemes are consistent with previous figures. **Top)** An abrupt termination of the RMCA is visible on the time-of-flight MIP. The corresponding RMCA vascular territory is highlighted in orange. **Middle)** A 2D projection of peak velocities throughout the Circle of Willis is displayed, with the peak velocity listed for each artery. A map of relative cerebrovascular reactivity (CVR) delays and their distributions within each vascular territory indicate longer delays in the RMCA territory. Relationships between median CVR delay and peak velocity for each vessel are plotted. **Bottom)** Flow values are reported next to each artery. A map of CVR amplitude and the distributions within each vascular territory indicate relatively preserved amplitude in the RMCA territory. Relationships between median CVR amplitude and average tissue perfusion for each vessel are plotted. Note: average tissue perfusion represents flow normalized to territory volume.

## Discussion

In a preliminary cohort spanning the adolescent age range, we assessed the normative relationship between macrovascular and microvascular cerebral hemodynamics using a novel combination of 4D flow MRI and BOLD cerebrovascular reactivity (CVR). We found that higher macrovascular velocities are associated with earlier relative CVR delays in downstream vascular territories. Conversely, higher macrovascular flows are associated with higher CVR amplitudes in downstream vessels, after accounting for differences in territory volumes. Both relationships change with age during adolescence. Similar trends were observed in moyamoya disease, emphasizing the importance of comprehensive cerebrovascular assessments in pathology. Further details regarding these findings are discussed below.

### Relationship between macrovascular peak velocity and tissue-level CVR delay

We observed that higher macrovascular peak velocities are associated with earlier relative CVR delays in corresponding vascular territories. In other words, brain regions supplied by arteries with higher blood velocity also respond earlier to a vasoactive stimulus (e.g., elevated CO_2_ from a breath-hold task). This was consistent with our hypothesis – an earlier response to a vasoactive stimulus may reflect shorter transit times of hypercapnic blood due to higher velocity in the supplying artery. However, CVR is a complex phenomenon that cannot be fully explained by velocities through intracranial arteries alone. CVR delays reflect the time it takes for hypercapnic blood to arrive at local brain regions (vascular transit time) and the time it takes for local arterioles to dilate in response to altered blood gas levels and to subsequently influence the venous-weighted BOLD contrast mechanism (vasodilatory dynamics).^53,56^ Variations in the vascular transit time may be due to higher velocities or shorter path lengths to the local region.^56,94^ It is important to characterize the contribution of vascular transit times and vasodilatory dynamics to CVR delay measurements, particularly in pathology where one or both may be affected.^56^ We cannot completely tease the two apart in our data, but our observation that resting macrovascular velocity and hypercapnic CVR delay are tightly coupled adds insight to this distinction. Future studies could incorporate a multi-post-labeling-delay ASL acquisition to investigate how arterial transit dynamics are related to macrovascular velocity and CVR delay. This information is particularly relevant as CVR delay becomes an increasingly popular metric in healthy individuals and clinical populations.^45^

While our study is the first to explore the relationship between macrovascular velocities and microvascular delays in healthy individuals, several studies have examined regional variations in blood velocity, arterial transit time, and CVR delay. Similar to our findings, Wu et al.^27^ reported highest 4D flow velocities in the MCAs, with lower velocities in the ACAs and PCAs. Arterial transit times measured with ASL follow a similar pattern in healthy adolescents, with significantly longer times in the PCA territory.^95^ Regional variations in BOLD CVR delays have also been widely reported in healthy individuals,^50,51,53,96–98^ with shortest delays (earliest responses) observed in the MCA territory and longest delays (latest responses) in the PCA territory.^96^

### Relationship between macrovascular flow and tissue-level CVR amplitude

As hypothesized, higher macrovascular blood flow is associated with higher CVR amplitudes in downstream vascular territories, after accounting for differences in vascular territory volumes. Although macrovascular flow is typically reported without normalizing to tissue volume (in units of ml/min), we implemented this normalization to facilitate comparisons between subjects with different brain sizes and to be consistent with previous literature. Most prior studies comparing macrovascular with microvascular flow utilize ASL or PET-derived tissue-level perfusion measurements, which reflect the volume of blood delivered per 100g of tissue per minute (ml/100g/min). When we do not account for territory volumes, the relationship between macrovascular flow and CVR amplitude is inverted, emphasizing the importance of this consideration to interpreting flow and CVR relationships. However, there are caveats to note in our comparison. First, computation of territory volumes included both gray matter and white matter, while median CVR was computed in gray matter. White matter CVR was excluded due to low BOLD SNR at 3T, leading to lower confidence in these CVR estimates. Additionally, we observed little variation in territory volumes across subjects, which may be a consequence of using an atlas derived from adults to delineate these regions (See “Limitations and Future Work”).

The positive association between flow and CVR amplitude is generally consistent with the small body of literature exploring similar relationships. Similar to this study, Clark et al.^64^ reported a strong positive relationship between mean arterial flow through the MCAs and ICAs (measured with 4D flow) and local tissue perfusion (measured with ASL) in cognitively healthy adults at risk for Alzheimer’s. In our previous work, we found a positive correlation between baseline tissue perfusion (measured with ASL) and BOLD CVR amplitude in healthy young adults, although both were tissue-level metrics.^82^ This paper^82^ provides a thorough commentary on other studies reporting positive correlations between baseline cerebral blood flow and BOLD CVR across individuals,^38,99,100^ and the few reports of *negative* correlations between tissue-level blood flow and CVR amplitude.^101,102^

Our findings provide evidence that CVR measured via BOLD fMRI is closely related to the inflow from upstream cerebral arteries. This is unsurprising, since large cerebral arteries are important determinants of microvascular pressure and perfusion,^63,64^ which in turn influence arteriolar dilation capacity.^65–67^ However, the relationship between macrovascular flow and microvascular CVR may not be causal and could instead be attributed to other factors such as differences in the size, density, and innervation of vessels supplying a given territory.^96^ To further probe this relationship, it would be helpful to modulate baseline macrovascular conditions within-subjects to test the subsequent CVR response, as done in previous adult studies.^65–67^ Regardless of the relationships we observe between such parameters, the degree of regional variation in macrovascular flow and CVR amplitude are consistent with previous studies. In healthy young individuals, on average, cerebral blood flow is distributed such that 21% goes through the MCAs, 12% through the ACAs, and 8% through the PCAs.^103^ Our flow measurements (before normalizing to tissue volume) are also within the range of previous 4D flow MRI reports.^104,105^

Lastly, few studies quantify regional heterogeneity of CVR amplitude in healthy individuals. Consistent with our findings, CVR is often highest in posterior brain regions supplied by the PCA.^92,96,98,106^ This may be due to microvascular anatomy in the occipital cortex, which contains a disproportionate number of blood vessels that could contribute to high BOLD signal amplitude.^96^ However, this could also be an artifact of higher SNR in the posterior cortex due to proximity of the head coil in the scanner.

### Symmetry of hemodynamic relationships

In healthy individuals, we observed symmetry in the hemodynamic relationships between hemispheres, but deviations from this pattern are expected in cerebrovascular pathologies which are often asymmetric. Although our hemispheric analysis is limited at the subject-level due to few data points, this approach could be used to glean insights across disease cohorts.

### Changing hemodynamic relationships in adolescence

Our exploratory analysis revealed age-related trends in the relationship between macrovascular and microvascular hemodynamics, though it is difficult to interpret the underlying mechanisms from our data. We observed opposite trends in the relationships for timing and flow parameters; the relationship between peak velocity and CVR delay decreases throughout adolescence, while the relationship between flow and CVR amplitude strengthens. To further probe these observations, we investigated how each macrovascular and microvascular parameter changes with age in the MCAs, ACAs, and PCAs, and their associated territories (combining measurements from left and right hemispheres).

We observed a significant decrease in peak velocity with age in all three vessels (Figure S2), which agrees with 4D flow^27^ and TCD studies^39,107^ demonstrating significant declines in flow velocity in major intracranial arteries, beginning around age 6 and continuing through adulthood. We also observed age-related trends in CVR delay, though this finding is difficult to interpret because the regional values are relative to a gray matter reference rather than absolute values. These trends are likely due in part to regional differences in path length based on the positive relationship we observed between CVR delay and gray matter volume in our cohort (Figure S4). The range of CVR delay values increased with gray matter volume, potentially reflecting higher variability in regional transit times in larger brains.

In contrast, we did not observe significant age-related changes in average tissue perfusion or CVR amplitude. Previous investigations of flow^103,108^ and CVR amplitude^38,109^ during adolescence have reported age-related changes, though most commonly these studies evaluate tissue-level perfusion rather than macrovascular flow. These reports also indicate a nonlinear modeling approach might be more appropriate for capturing age-related perfusion and CVR dynamics, which our sample size is not sufficient to characterize.

While we cannot make definitive conclusions about the underlying mechanisms, relationships between macrovascular and microvascular hemodynamics are changing between childhood and adulthood. This likely reflects the series of structural and functional changes the brain undergoes during this period to support increasingly complex behavior. The developmental trajectories of gray matter density^32^ and volume^110^ in the brain have been well-documented, but cerebrovascular maturation is not well understood.^40^ Our findings emphasize the need for more sophisticated cerebrovascular imaging assessments in adolescents to better characterize these developmental changes.

### Implications for understanding pathological mechanisms in complex patients

Our moyamoya arteriopathy case study underscores the importance of comprehensive cerebrovascular assessments, encompassing both macro- and micro-vascular circulation, as well as flow and timing parameters, to understand complex pathology. Despite compromised blood flow in this individual’s right MCA leading to prolonged CVR delays, CVR amplitude in the affected MCA territory is relatively preserved. This is likely due to compensatory flow patterns through the Circle of Willis and the development of collateral vessels that maintain sufficient blood supply to the region.^111^ Thus, while a macrovascular assessment might raise alarm, understanding how this stenosis translates to tissue-level hemodynamics may be more informative for assessing stroke risk and guiding aggressive treatment options like revascularization surgery.

Combining macrovascular and microvascular MRI measurements may be particularly impactful in sickle cell disease, a condition often associated with moyamoya. Cerebrovascular complications in sickle cell disease include large artery stenosis, which may cause symptomatic ischemic stroke, as well as more subtle tissue-level “silent” infarcts. As mentioned in the introduction, transcranial Doppler ultrasound (TCD) is the current clinical standard for stroke screening in sickle cell disease,^16^ but TCD measurements are operator-dependent and offer limited spatial coverage of intracranial vessels.^112^ TCD also requires an accessible acoustic window, which becomes difficult as the skull ossifies throughout childhood. As a result, many adolescents with sickle cell disease stop receiving examinations around the age of 16, despite continued stroke risk. 4D flow MRI could address these limitations and serve as a surrogate for TCD by providing blood velocity measurements in all major cerebral arteries in one acquisition, in both children and adults with sickle cell disease. However, blood velocities are not associated with tissue-level perfusion^18^ or with silent infarct occurrence^19,20^ in children with sickle cell disease, which suggests silent infarct pathology might be distinct from macrovascular processes assessed by TCD. Thus, a complementary biomarker of microvascular health, such as CVR, may be more appropriate for predicting the occurrence of silent infarcts and has already been implemented at some children’s hospitals. Recent clinical guidelines^113^ recommend MRI/MRA brain exams for children and adults with sickle cell disease, although these exams only provide anatomical data. Our study suggests that specialized MRI techniques like 4D flow and CVR could be added to provide more comprehensive information regarding macro- and micro-vascular physiology.

### Considerations for clinical application

Although 4D flow MRI and BOLD CVR offer valuable insights into cerebrovascular health, there are important caveats for clinical application. The long 4D flow acquisition time is currently impractical for routine clinical use but can performed in under 6 minutes with additional acceleration strategies^114^ and machine learning techniques.^115^ Breath-hold CVR has been successfully implemented in pediatric cohorts^60^, but alternative approaches for the breathing task^116^ or post-processing^91^ may be necessary to address compliance challenges. As with any MRI acquisition, these methods are more expensive than alternatives and susceptible to motion artifacts. Mitigating these challenges will be crucial.

At this preliminary stage, we see this imaging protocol being more useful for mapping regional hemodynamics on an individual basis, rather than determining cut-off values for stratification. For example, within-subject information could be used to identify regions with atypical CVR characteristics based on comparisons to homologous regions in the opposite hemisphere. Changes in these within-subject relationships could be monitored over time. Furthermore, we do not necessarily intend for these methods to replace clinical protocols, but rather to serve as tools for understanding complex hemodynamic mechanisms in neuropathophysiology and potentially only in research settings. Depending on the relationships between macrovascular and microvascular hemodynamics in target patient cohorts, a breath-hold CVR scan could offer spatially sensitive detection of both macrovascular and microvascular pathology. Alternatively, combining both scans may be necessary to distinguish between macrovascular and microvascular circulation effects. Finally, we recognize other MRI approaches can provide similar information, and our main call to researchers and physicians is to consider combining and relating macrovascular and microvascular assessments for a more holistic approach.

### Limitations and future work

We acknowledge our study is limited by a small sample size. This normative dataset should be expanded to confirm the observed relationships and to probe age-related changes with more complex modeling approaches.^109,110^ A larger, multi-session study could address the high inter-subject variability in hemodynamic relationships we observed. A test-retest analysis would be useful to evaluate true measurement variability relative to individual, age-dependent variability.

Additionally, we only examined macrovascular-microvascular relationships in one moyamoya vasculopathy case, but more work is necessary to characterize these relationships in pathology. Future investigations to elucidate the clinical potential of this imaging approach should include longitudinal studies to track changes in hemodynamic metrics over time, correlate them with clinical outcomes, and compare them to current standards of care.

Our study was also limited by using an atlas specifically developed for healthy adults to define vascular territory regions.^83,84^ These atlas-defined territories are likely less generalizable to younger participants and may be inaccurate for the moyamoya subject, where collateral flow pathways may alter the boundaries of the brain region supplied by each vessel. Despite these limitations, an atlas approach offers ease of implementation and improved generalizability. The impact on our conclusions should be minimal due to the large number of voxels included in CVR calculation. Additionally, in the moyamoya case, we found that changing the boundaries of the affected RMCA territory did not substantially alter CVR estimates (Figure S3). A pediatric-specific territory atlas does not exist, so alternative solutions include manual delineation^117^ or velocity-selective ASL^118^. The former approach is labor-intensive and would require neuroradiologist verification. Velocity-selective ASL acquisition would prolong an already lengthy protocol and suffers from low SNR, motion sensitivity, and contamination from CSF signals.^119^

Overall, our study provides unique insight into macrovascular and microvascular hemodynamics during adolescence and important normative data for future investigations in pediatric cerebrovascular pathology. The combination of 4D flow MRI and BOLD CVR provides a framework for quantitative, multi-scale evaluation of cerebral hemodynamics. Future work should apply a similar multi-scale framework in both normative and pathological cohorts to expand our understanding of cerebrovascular maturation and to address critical challenges in adolescent cerebrovascular imaging assessments.

## Acknowledgements

This work was supported by Northwestern University’s Center for Translational Imaging. Our sincere gratitude goes to Rachael Young for her advice on scan protocols and support with data acquisition. Thank you to Kai Yang and Adam Richter for their contributions to the 4D flow parametric mapping code. We are grateful to the participants and their families for making this research possible.

## CRediT Author Contribution Statement

**Kristina Zvolanek:** Conceptualization, Methodology, Software, Formal analysis, Investigation, Writing – Original Draft, Writing – Review & Editing, Visualization, Project administration, Funding acquisition. **Jackson Moore:** Methodology, Software, Validation, Writing – Review & Editing. **Kelly Jarvis:** Methodology, Software, Writing – Review & Editing. **Sarah Moum:** Conceptualization, Writing – Review & Editing, Supervision. **Molly Bright:** Conceptualization, Methodology, Resources, Writing – Review & Editing, Supervision, Project administration, Funding acquisition.

## Supplementary material

Supplementary material for this paper can be found at http://jcbfm.sagepub.com/content/by/supplemental-data.

## Footnotes

a A manuscript describing this analysis method is currently under review. We plan to add this citation later in the review process.

