## Supplemental Material for "Macrovascular blood flow and microvascular cerebrovascular reactivity are regionally coupled in adolescence"

### Methods

***Table S1. Magnetic resonance imaging (MRI) scan parameters***

| | **Time-of-flight** | **Dual-*venc* 4D flow** | **Multi-echo BOLD fMRI** | **T1-weighted MPRAGE** |
|---|---|---|---|---|
| Plane | Axial | Axial | Axial | Sagittal |
| TR (ms) | 21.0 | 5.9 | 1500 | 2170 |
| TE (ms) | 3.42 | 3.3 | 10.6/27.83/45.06/ 62.29/79.52 | 1.69/3.55/5.4 1 |
| Flip angle (°) | 18 | 15 | 70 | 7 |
| In-plane resolution ($mm^{2)}$ | 0.3 x 0.3 | 1 x 1 | 2.5 x 2.5 | 1 x 1 |
| Slice thickness (mm) | 0.5 | 1 | 2.5 | 1 |
| Number of slices | 4 slabs of 40 slices | 44 | 52 | 176 |
| Field-of-view (mm) | 182 x 200 | 192 x 220 | 210 x 210 | 256 x 256 |
| Bandwidth (Hz/Px) | 186 | 460 | 2480 | 650 |
| Acceleration factor | GRAPPA = 2 | k-t GRAPPA R = 5 | Multiband = 4; GRAPPA = 2 | GRAPPA = 2 |
| Scan time | 5 min 33 s | 10-12 minutes | Depends on task length | 5 min 12 s |
| Other | -- | low-*venc* = 0.8 m/s; high-*venc* = 1.6 m/s; 82.6 ms temporal resolution | -- | TI = 1160 ms |

#### *Functional MRI Pre-processing*

The T1-weighted MPRAGE was processed with *fsl_anat,* which involved bias-field correction, brain extraction, and segmentation of gray matter, white matter, and cerebrospinal fluid tissues. Linear registration was performed with FSL *flirt* to co-register the T1-weighted image to the brain-extracted SBRef image of the first echo. Non-linear registration was performed with FSL *fnirt* to co-register both the functional (first echo SBRef) and anatomical images to the MNI152 $6^{th}$ generation template (FSL version, 2 mm resolution).[1]

Volume realignment of the functional data was performed using the SBRef image of the first echo as the reference and applying the spatial transformation to all subsequent echoes.[2,3] An optimal combination of the five echoes was created with *tedana.*[4,5] The pair of first echo SBRef images with reverse phase-encoding directions was used to perform field distortion correction with FSL *topup.*[6] Then, the optimally combined, distortion-corrected data were brain-extracted with FSL *bet* and used as the input for CVR modeling.

#### *CVR Amplitude and Delay Estimation*

First, 61 shifted variants of the demeaned $P_{ET}CO_2$ regressor were created in 0.3 s increments, ranging ±9 s from an initial "bulk" shift.[7] An extended delay range (±15 s) was used for the individual with moyamoya. fMRI data were modelled by a design matrix consisting of the shifted $P_{ET}CO_2$ signal, Legendre polynomials, and six motion parameters. Each lagged-GLM was fitted via orthogonal least squares and the maximum full model $R^2$ was identified for each voxel.[7] The corresponding shift (in seconds) determined CVR delay, and the associated beta coefficient was extracted and rescaled to be expressed in percentage BOLD signal change (%BOLD).

#### *Impact of Right MCA Boundaries in Moyamoya Case Study*

To test whether changing the boundaries of the affected right MCA (RMCA) territory impacts the resulting CVR values, we created 6 new versions of the RMCA mask by dilating or eroding the arterial territory atlas mask by 3x3x3, 6x6x6, or 9x9x9 voxels, respectively. We then computed the median CVR amplitude and delay within these masks to compare to the value obtained with the original atlas (Figure S3).

### Results

#### Table S2. Linear mixed effects model results for hemispheric relationships between CVR delay and peak velocity

| | L Hemisphere CVR Delay | | R Hemisphere CVR Delay | | Both Hemispheres CVR Delay | |
|---|---|---|---|---|---|---|
| **Fixed Effect: Peak Velocity** | Estimate ± SE | t-statistic | Estimate ± SE | t-statistic | Estimate ± SE | t-statistic |
| Slope | -0.912 ± 0.380 | -2.40* | -1.268 ± 0.402 | -3.16** | -1.055 ± 0.361 | -2.93* |
| Intercept | -0.201 ± 0.075 | -2.67** | -0.051 ± 0.068 | -0.74 | -0.122 ± 0.064 | -1.89** |
| **Random Effect: Subject** | Variance | | Variance | | Variance | |
| Slope | 0.456 | | 0.463 | | 0.727 | |
| Intercept | 0.187 | | 0.113 | | 0.033 | |
| Residual | 0.313 | | 0.360 | | 0.103 | |

Note: *p < 0.05, **p < 0.01, ***p < 0.001

#### Table S3. Linear mixed effects model results for hemispheric relationships between CVR amplitude and tissue perfusion

| | L Hemisphere CVR Amplitude | | R Hemisphere CVR Amplitude | | Both Hemispheres CVR Amplitude | |
|---|---|---|---|---|---|---|
| **Fixed Effect: Avg. Tissue Perfusion** | Estimate ± SE | t-statistic | Estimate ± SE | t-statistic | Estimate ± SE | t-statistic |
| Slope | 0.001 ± 0.0003 | 3.04** | 0.001 ± 0.0003 | 4.43*** | 0.001 ± 0.0003 | 3.86*** |
| Intercept | 0.382 ± 0.026 | 14.60*** | 0.388 ± 0.025 | 15.81*** | 0.388 ± 0.025 | 14.75*** |
| **Random Effect: Subject** | Variance | | Variance | | Variance | |
| Slope | 0.001 | | 0.001 | | 0.001 | |
| Intercept | 0.089 | | 0.083 | | 0.089 | |
| Residual | 0.031 | | 0.029 | | 0.027 | |

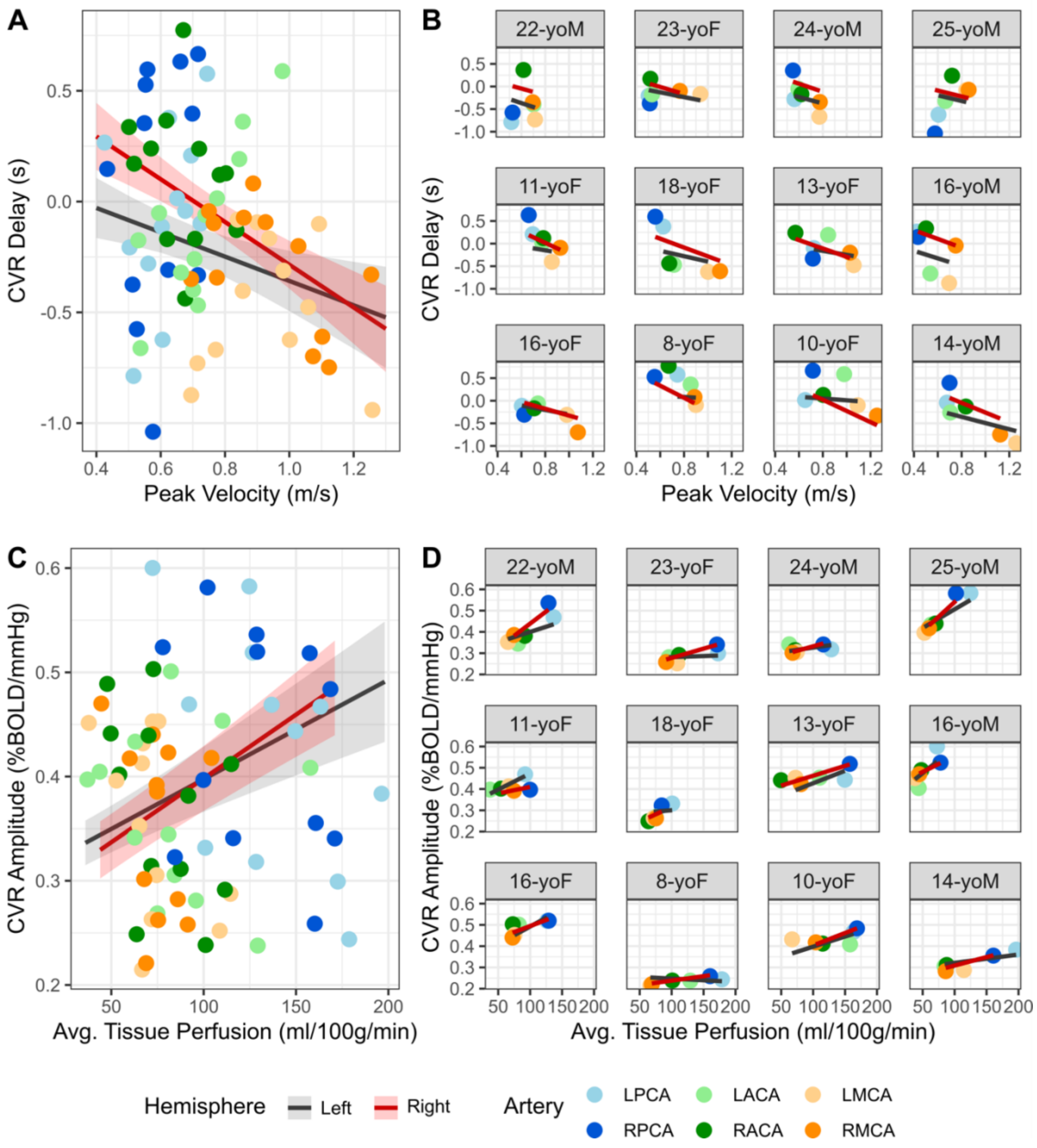


***Figure S1.*** *Hemispheric relationships between macrovascular and microvascular hemodynamics. Gray lines indicate left hemisphere trends; red lines indicate right hemisphere trends. Peak velocity and CVR delay relationships for* ***A)*** *the group and* ***B)*** *each subject. Average tissue perfusion and CVR amplitude relationships for* ***C)*** *the group and* ***D)*** *each subject. Hemispheric linear fits were plotted using estimates from the linear mixed effects models detailed in Tables S2 and S3.*

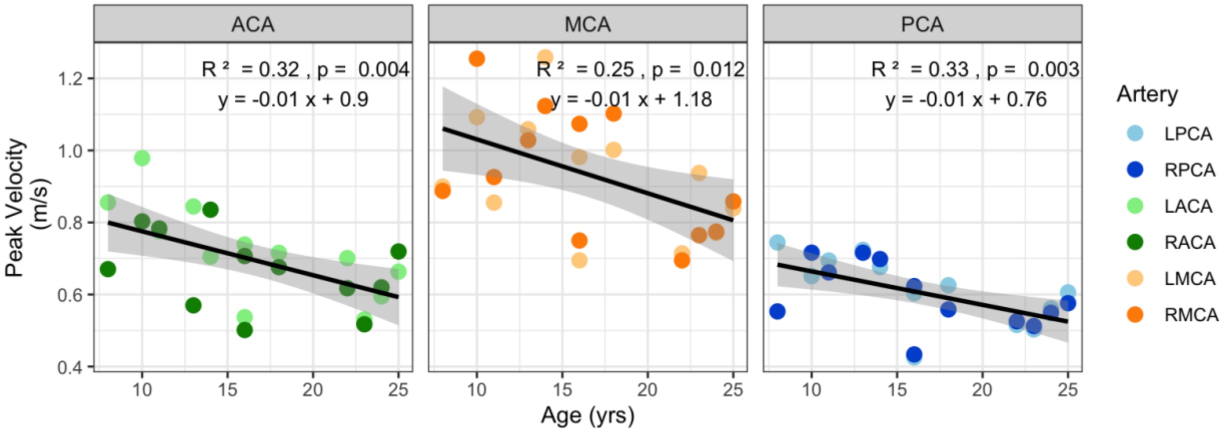


***Figure S2.*** *Age-related changes in peak velocity for the anterior cerebral arteries (ACA), middle cerebral arteries (MCA), and posterior cerebral arteries (PCA). Values from the right and left hemispheres were combined to evaluate the trend for each vessel, though they are indicated in the plots by different colors. The least-squares fit line, corresponding equation, and associated* $R^2$ *and p-values are plotted for each vessel.*

#### *Impact of Right MCA Boundaries in Moyamoya Case Study*

Figure S3 shows how changing the RMCA boundaries affects CVR estimates in the participant with moyamoya arteriopathy. CVR amplitudes are consistent regardless of the RMCA mask boundaries, ranging from 0.20 to 0.22 %BOLD/mmHg. There is a wider range of CVR delays (2.31 to 3.63 seconds), with larger differences in more dilated masks. These results are in line with the CVR maps shown in Figure 6. While CVR amplitude is preserved across the brain, there is a focal region of increased CVR delay. It makes sense then, that when dilating the mask to include "healthy" tissue in the RACA or RPCA territories, we see a decrease in the median delay value.

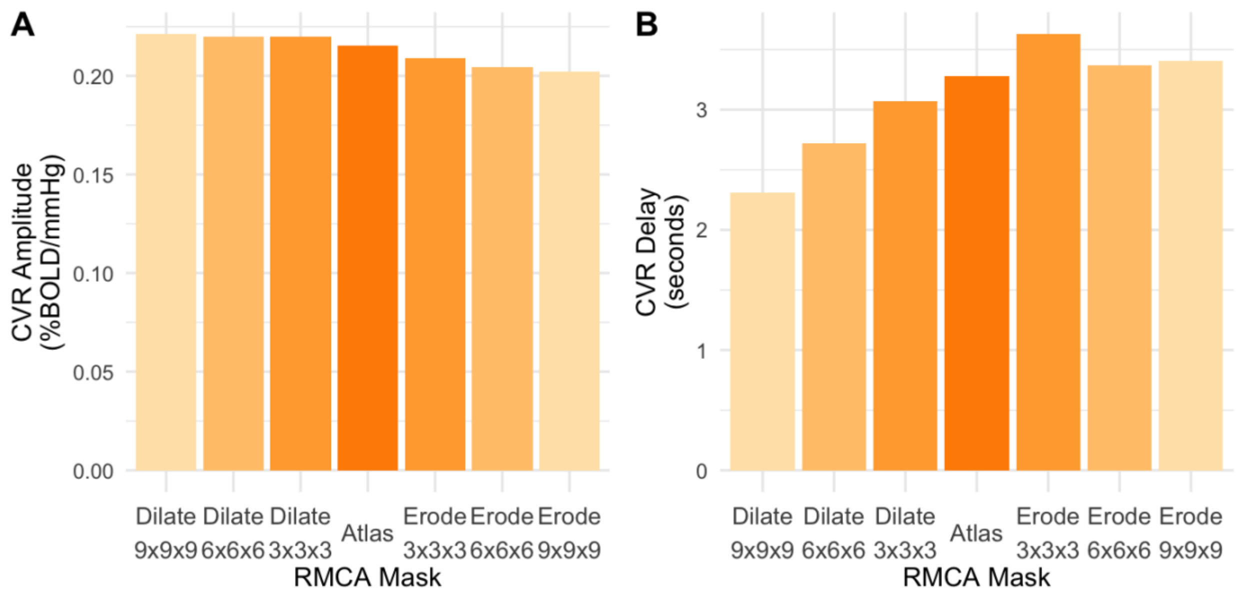


***Figure S3.*** *Median cerebrovascular reactivity (CVR)* ***A)*** *amplitude and* ***B)*** *delay in different right middle cerebral artery (RMCA) masks for the participant with RMCA moyamoya arteriopathy. The RMCA mask from a vascular territory atlas is indicated in dark orange. This mask was then successively dilated or eroded in increments of 3x3x3 voxels. Medians reflect only voxels in gray matter, consistent with analysis in the main text.*

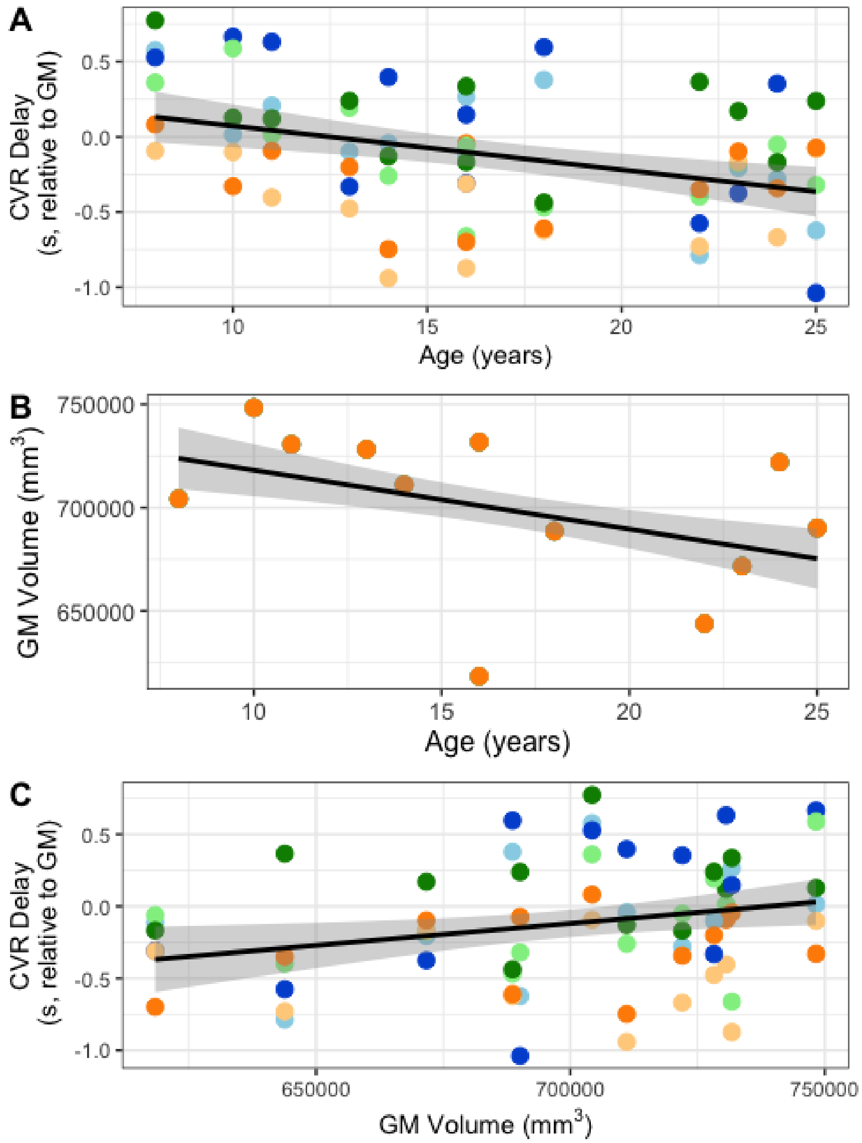


***Figure S4.*** *Age-related changes in cerebrovascular reactivity (CVR) delay are mediated by whole-brain gray matter (GM) volume. A) Changes in median CVR delay with age. Points represent the median value in one of six vascular territories (left and right ACA, MCA, PCA) for each subject. The color scheme is consistent with other figures in the manuscript. B) Age-related changes in gray matter volume. C) Changes in median CVR delay with gray matter volume.*
